# Chromatin conformation remains stable upon extensive transcriptional changes driven by heat shock

**DOI:** 10.1101/527838

**Authors:** Judhajeet Ray, Paul R. Munn, Anniina Vihervaara, Abdullah Ozer, Charles G. Danko, John T. Lis

## Abstract

Heat shock (HS) initiates rapid, extensive, and evolutionarily conserved changes in transcription that are accompanied by chromatin decondensation and nucleosome loss at heat shock loci. Here we have employed *in situ* Hi-C to determine how heat stress affects long-range chromatin conformation in human and *Drosophila* cells. We found that compartments, topologically-associated domains (TADs), and looping interactions all remain unchanged by an acute HS. Knockdown of Heat Shock Factor 1 (HSF1), the master transcriptional regulator of the HS response, identified HSF1-dependent genes and revealed that up-regulation is often mediated by distal HSF1 bound enhancers. HSF1-dependent genes were usually found in the same TAD as the nearest HSF1 binding site. However, the HSF1 binding sites and the target promoters did not exhibit a focal increase in contact frequencies compared with surrounding regions, nor did we find evidence of increased contact frequency following HS. Integrating information about HSF1 binding strength, RNA polymerase abundance at the enhancer, and contact frequency with a target promoter in the non-heat shock (NHS) condition accurately predicted which up-regulated genes were direct targets of HSF1 during HS. Our results suggest that the chromatin conformation necessary for a robust HS response is pre-established in normal (uninduced) cells of diverse metazoan species.

## INTRODUCTION

Proximal and distal regulatory elements coordinate cell type specific transcriptional programs necessary for normal cellular function. HS response is a well-studied model system for understanding gene regulation in metazoan organisms, including flies (1–5), mice (6) and humans (7, 8) causing both up-regulation of hundreds and down-regulation of thousands of target genes. HS induces binding of HSF1 to numerous heat shock elements (HSEs) across the genome. HSF1 binding to HSEs increases the rate at which paused RNA polymerase II (Pol II) is released into productive elongation at up-regulated genes (6, 8). Although the majority of HSF1-activated genes have promoter-bound HSF1, many do not (6, 7, 9). This indicates that HSF1 can regulate gene transcription through distant enhancer interactions.

The three-dimensional structure of chromatin in the nucleus is proposed to play a fundamental role in gene regulation by facilitating or restricting regulatory element interactions. Gene activation during HS is linked to dramatic changes in chromatin. Loci encoding activated genes form highly-visible puffs in polytene chromosomes of *Drosophila* upon HS (10, 11), and biochemical assays reveal massive changes in nuclease sensitivity and nucleosome loss in non-polytene cells (12, 13). Dramatic transient changes in histone modification and chromatin composition also occur by the recruitment of specific transcription factors, chromatin remodelers and histone modifiers (8, 14, 15). The extent of these changes along the chromosome and how they might influence long-range interactions between distal DNA sequences measured by Hi-C remains unclear.

In this study, we have used *in situ* Hi-C (16) to map the genomic contacts in human K562 and *Drosophila* S2 cells subjected to heat shock. We observed no evidence for global changes in compartments, TADs, or looping interactions in heat shocked cells. Despite a lack of changes in Hi-C data, integrating information about HSF1 binding strength and contact frequency with a target promoter accurately predicted which up-regulated genes were direct HSF1 targets. Thus, we propose that chromatin architecture necessary for HS response is pre-established in both human and *Drosophila* cells, potentially reflecting an evolutionarily conserved mechanism that enables cells to respond rapidly to stress.

## RESULTS

### Global chromatin architecture is conserved during HS despite dramatic transcriptional changes

We have previously reported that HS induces transcriptional changes in thousands of genes in humans, mice, and *Drosophila* (5, 6, 8). To understand how changes in transcription correlate with changes in three-dimensional chromatin architecture, we performed in situ Hi-C (16), both before and after 30 min of HS in the human chronic myelogenous leukemia K562 cell line (**Fig. 1A**). Hi-C libraries were sequenced to an estimated resolution of 10 kb in each condition (**Table S1**). We confirmed that biological replicates were highly correlated (Stratum-adjusted Correlation Coefficient [SCC] > 0.94) at 10 kb resolution using HiCRep (17), a method of correlating Hi-C data that compensates for distance dependence and domain structure (**Table S2**). Comparison between HS and NHS Hi-C contact maps revealed a highly similar distribution of contact pairs across the genome (**Fig. 1B**). Genome-wide analysis using HiCRep revealed that heat maps from HS and NHS were correlated to the same extent as biological replicates (**Table S2**). Thus, to a first approximation, we observed no evidence for differences in Hi-C contact maps between the two conditions.

**Figure 1:**
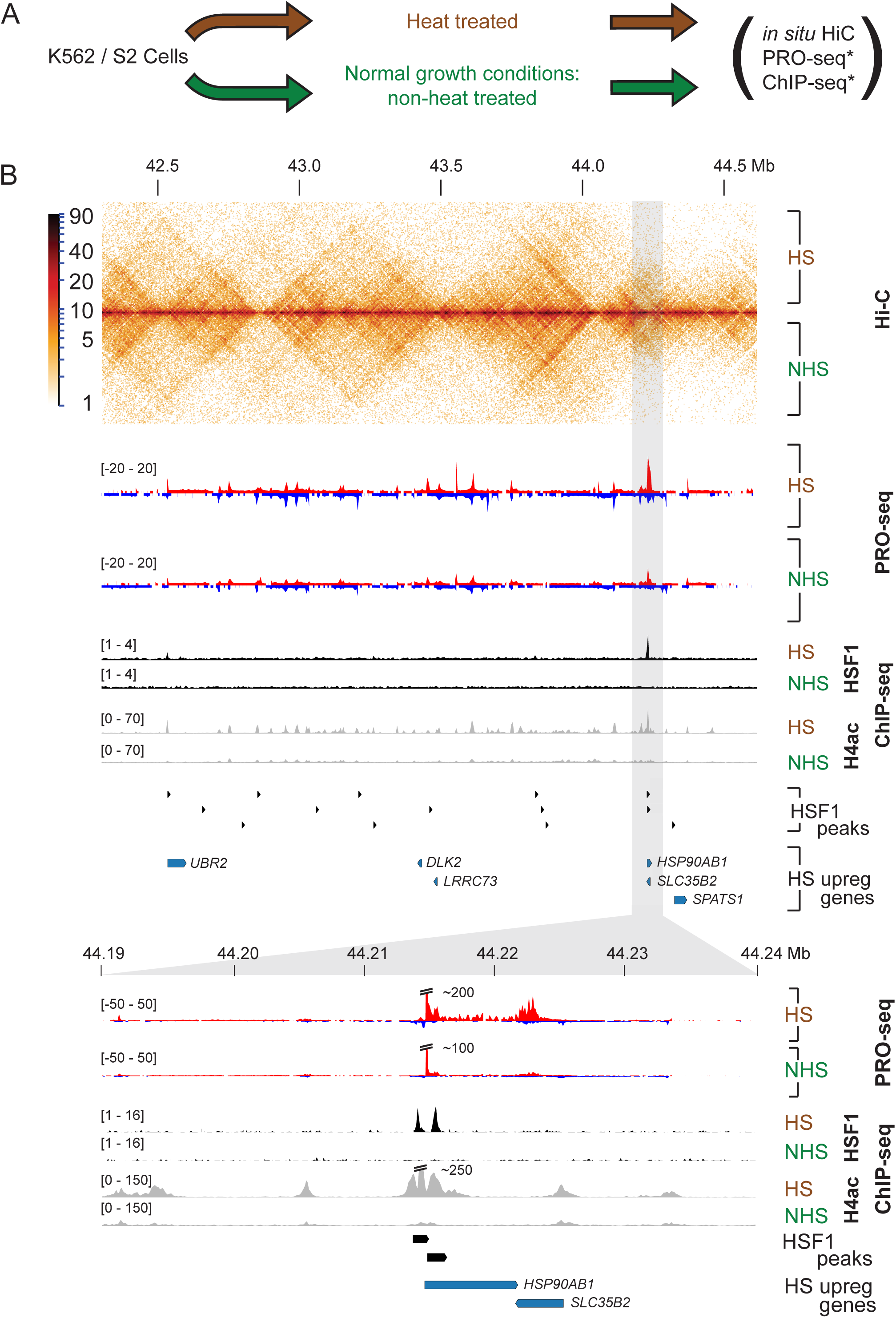
Global chromatin architecture is unaltered during HS induced transcriptional changes. A) Schematic of data sets generated and analyzed in this study. Human K562 or *Drosophila* S2 HS and NHS cells were used to generate in situ Hi-C data, and these datasets were compared to corresponding PRO-seq (5, 8) and ChIP-seq (4, 7, 8) data sets denoted in asterisk. B) Comparison of PRO-seq, *in situ* HiC, and ChIP-seq assays performed on heat shocked and non-heat shocked K562 cells. Grey region highlights a classical heat shock locus containing HSP90AB1 gene.

To determine whether HS changed chromatin conformation near HS regulated genes, we first classified genes as HS upregulated, downregulated, or unregulated using PRO-seq data (8). As not all up-regulated genes depend on HSF1 (5, 6), we used PRO-seq data from control and HSF1 RNAi knockdown in K562 cells to classify up-regulated genes into HSF1-dependent or -independent categories (Vihervaara et al. manuscript in prep). We classified 5,746 genes as unregulated with no detectable change in expression by PRO-seq, 227 genes as HSF1-dependent, 360 as upregulated but HSF1-independent, while 4,002 genes as downregulated (**Fig. S1**).

Examination of Hi-C heatmaps near regions with HSF1 up- or down-regulated genes revealed similar patterns between HS and NHS. For instance, a closer examination of a locus harboring a classical HS gene, HSP90AB1, showed HS enriched HSF1 binding near the promoter, 4.3-fold transcriptional activation, and a dramatic increase in acetylation of histone 4. However, HS did not cause any obvious change in the composition of Hi-C heatmap (**Fig. 1B**). Thus, variation between NHS and HS conditions were relatively minor at the whole chromosome scale.

### Chromosomal compartmentalization is preserved upon HS

Hi-C has revealed the presence of active (A) and inactive (B) compartments, corresponding to regions of open, transcriptionally active chromatin, and closed, silent chromatin domains, respectively, in mammalian genomes (16, 18). A subset of compartments switch between active (A) and inactive (B) states in a manner that correlates with gene expression changes (19). To investigate whether short durations of HS change compartment organization, we identified compartments at 40 kb resolution in HS and NHS data. Examination of compartments in Juicebox (16) revealed that compartments were highly correlated between the NHS and the HS conditions in K562 cells (**Fig. 2A**). Quantitative analysis of principal compartment scores revealed a high correlation in compartment strength genome-wide between NHS and HS conditions (Pearson’s R = 0.94).

**Figure 2:**
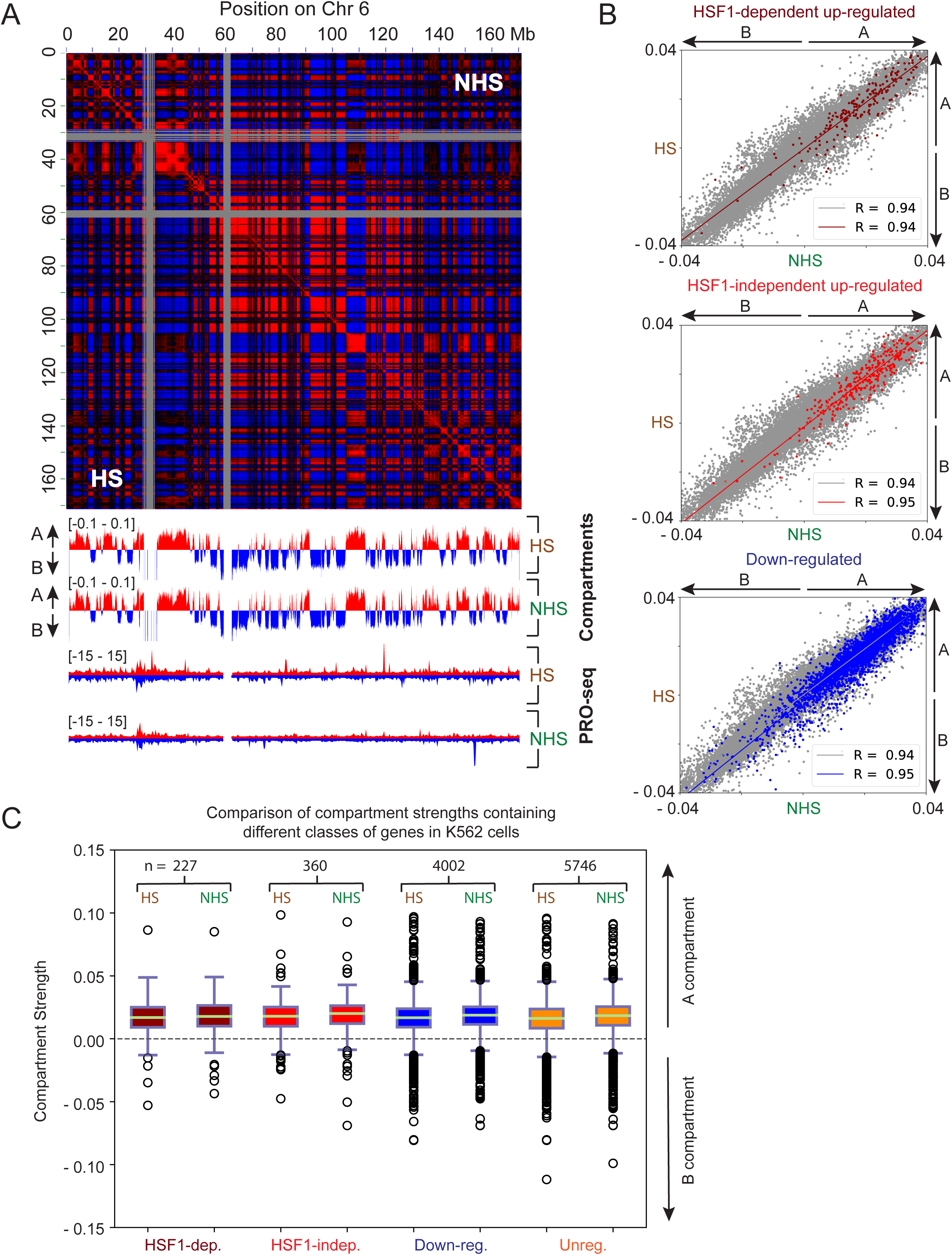
Chromosomal compartmentalization is preserved upon HS. A) Pearson correlation heatmap. Tracks below show first principal component (for both NHS and HS in K562 cells) - used to call compartments (shown here for bin sizes of 50Kb). B) Scatter plots show correlation between the strength of A and B compartment calls before and after HS. Grey - All compartments; Dark red - Compartments containing HSF1-dependent up-regulated genes (top); Red - Compartments containing HSF1-independent up-regulated genes (middle); Blue - Compartments containing down-regulated genes (bottom). R = Pearson’s correlation coefficient. Solid line in each plot represents the best fit line (linear regression). C) Boxplots showing the relative strength of compartments containing different classes of genes, for both HS and NHS conditions. Dark red - HSF1-dependent up-regulated genes; Red – HSF1-independent up-regulated genes; Blue - Down-regulated genes; Orange - Unregulated genes. n represents number of genes in each class.

If changes in transcription were accompanied by changes in chromatin compartmentalization, this would be most evident in compartments harboring HS regulated genes. However, compartment calls for each category were highly correlated between the NHS and HS conditions with a Pearson correlation coefficient ranging from 0.94 - 0.95, suggesting no major difference in compartmentalization upon HS for any of these categories of genes (**Fig. 2B**). The small differences observed were similar in magnitude to those observed between biological replicates from the same HS conditions (R = 0.92 – 0.93) (**Fig. S2**), suggesting that these differences are technical noise in the data rather than real biological signal. Both HS up- and down-regulated genes were predominantly localized in the active (A) compartment in both the NHS or HS conditions (**Fig. 2C**), consistent with previous observations of significant transcription of these genes before and after thermal stress (8). We observed small decreases in compartment strength for all classes of genes after HS. However, these changes were not correlated with the changes observed in transcription, indicating a transcription-independent mechanism. Taken together, these results show that compartmentalization remains largely unchanged during the first 30 min of HS.

### Heat shock does not affect TAD boundaries

A previous study found evidence that TAD boundaries change following a short duration of heat stress in *Drosophila* (20). To determine whether TAD boundaries change in our Hi-C data, we determined the insulation score (21) as a measure of TAD boundary strength in NHS and HS conditions. We found that insulation scores were highly similar between HS and NHS conditions (Pearson’s R = 0.98), suggesting that TAD boundaries remain stable in response to 30 min of HS (**Fig. 3A, 3B, and S3**).

**Figure 3:**
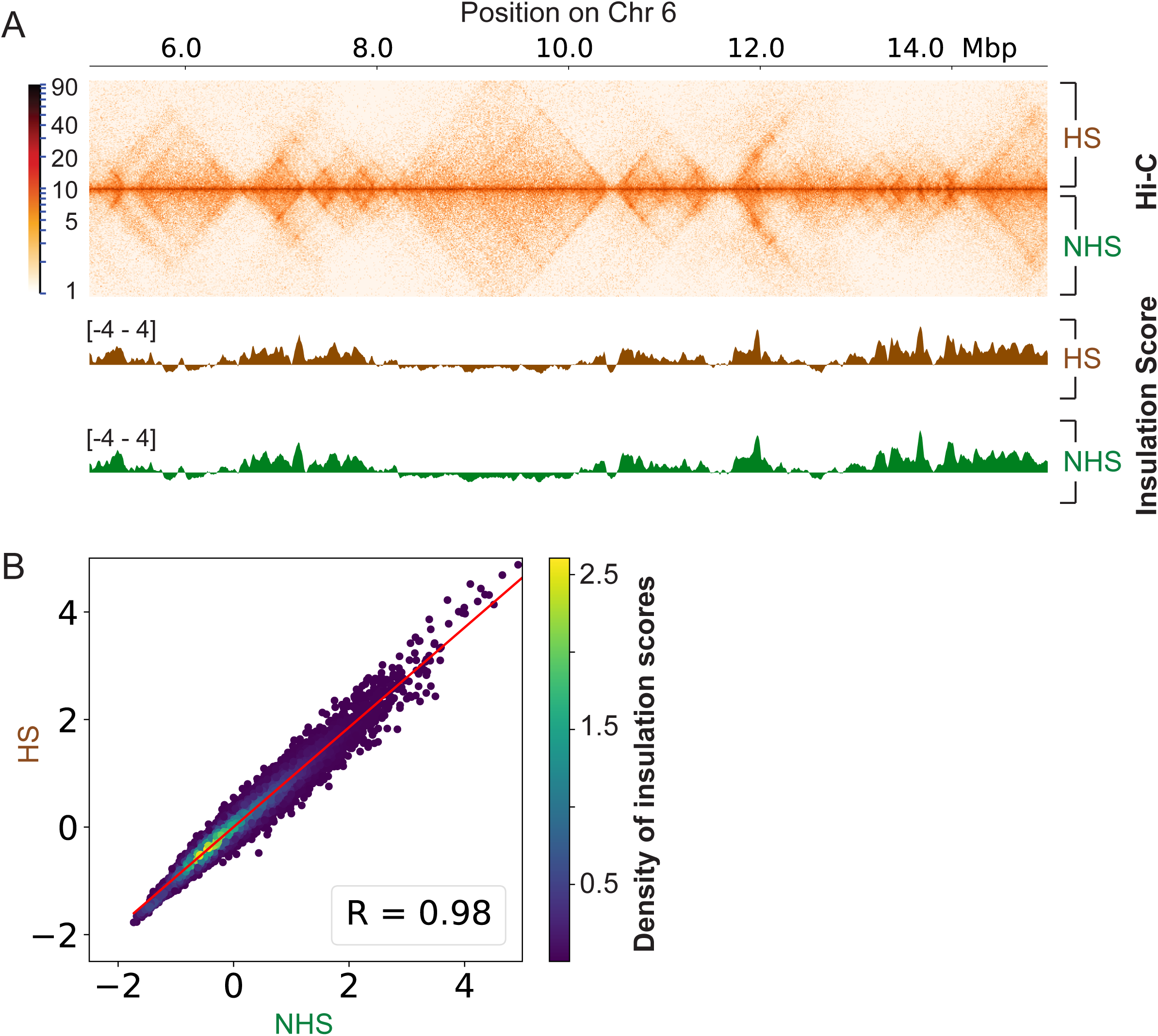
Heat shock does not affect TAD boundaries. A) 10 Mb region of chromosome 6 (in K562) showing comparison of HS and NHS Hi-C contact matrices (top). Comparison of insulation scores for same region, for both the conditions (bottom). B) Correlation of insulation scores for HS vs NHS conditions, for all chromosomes in K562. R = Pearson’s correlation coefficient. Red line represents the best fit line (linear regression).

### Heat shock does not affect enhancer-promoter contact frequencies

We asked whether HS induces changes in contact frequencies between enhancers and their target promoters. We focused on HSF1-dependent genes up-regulated, for which the failure of the gene to activate following HSF1 knockdown provided functional evidence of an interaction. The majority of HSF1-dependent genes had an HSF1 binding site located within 1.5 kb of the transcription start site (TSS) (**Fig. S4**). The distribution of contact frequencies between the promoter and the HSF1 bound peaks stratified by genomic distance revealed no significant increase in contact frequencies regardless of the distance between them (**Fig. 4A**).

**Figure 4:**
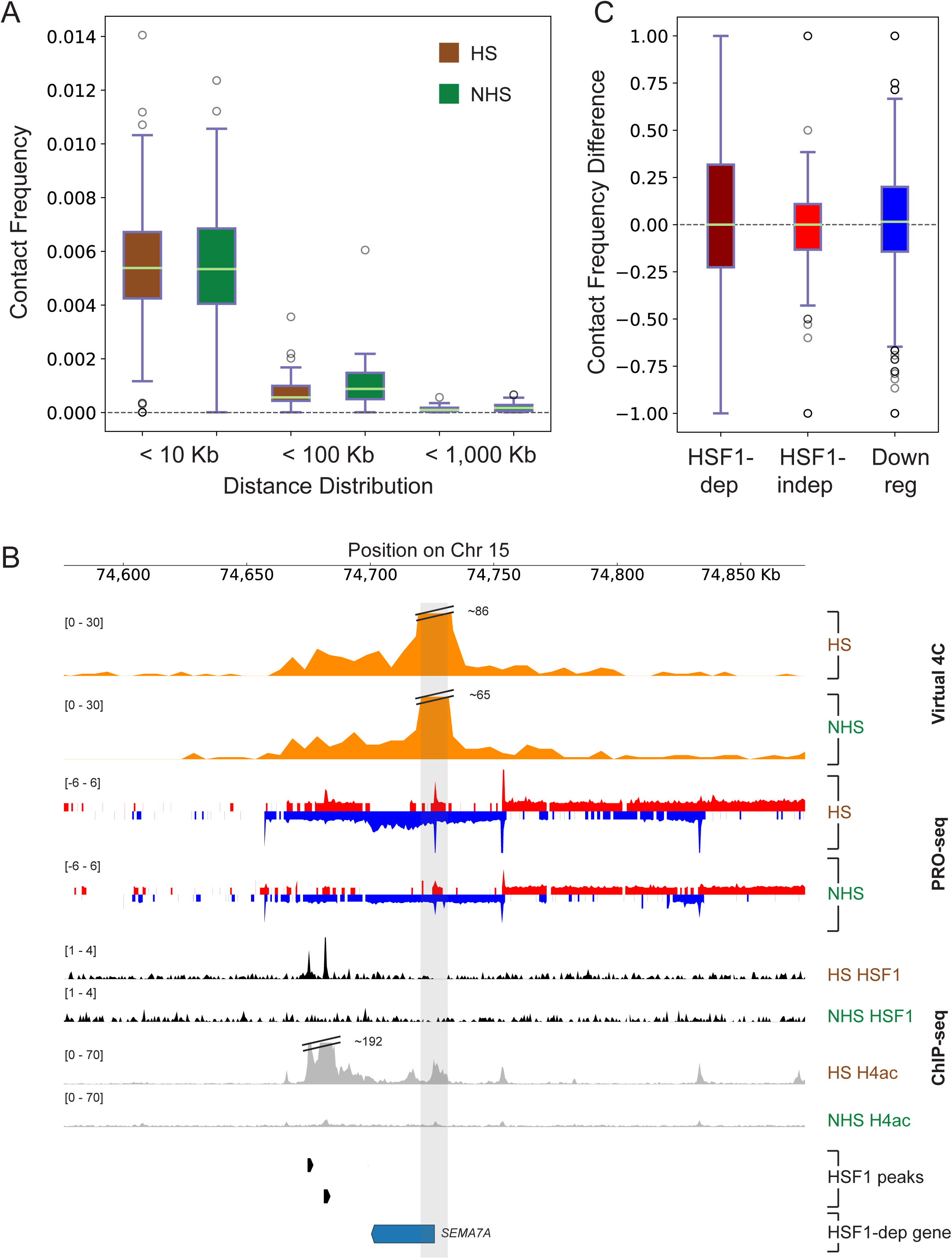
Comparison of contact frequencies between HSF1-dependent gene promoters and HSF1 binding sites under NHS and HS conditions. A) Distribution of contact frequency relative to linear distance of contact end points. Green line represents the median, boxes represent 25-75 percentile and whiskers represent 5-95 percentile. B) Virtual 4C plot showing contacts between *SEMA7A* TSS and its nearest HSF1 binding sites. PRO-seq tracks are included to show changes in transcription for this gene upon HS. ChIP-seq tracks of HSF1 and H4ac show HSF1 binding and accompanying active chromatin status respectively upon HS. C) Difference in contact frequencies between gene TSSs and distal (≥ 10 kb) enhancer sites for HSF1-dependent and HSF1-independent up-regulated, and HS down-regulated genes under NHS and HS conditions. Contact frequencies were calculated for HSF1-dependent genes between TSS and HSF1 binding sites; for HSF1-independent genes between TSS and active, up-regulated transcriptional regulatory elements; for down-regulated genes between TSS and active, down-regulated transcriptional regulatory elements.

This analysis was driven to a large extent by HSF1-dependent genes with an HSF1 binding site located near the promoter, and may therefore miss changes in contact frequency at genes regulated by distal HSF1 binding sites. We found many clear examples in which distal HSF1 binding sites played a role in gene activation at HSF1-dependent up-regulated genes. For example, *SEMA7A* did not show HSF1 occupancy at the promoter, but harbors two HSF1 binding sites at an enhancer cluster that show HS-dependent enhancer RNA (eRNA) transcription and reside ~35 kb downstream of the gene’s TSS (**Fig. 4B**). To quantitatively determine the connectivity of *SEMA7A* before and after HS, we surveyed contact frequencies between the *SEMA7A* promoter and all DNA within 100 kb up- and downstream using a virtual 4C analysis (**Fig. 4B**). We found that both HSF1 binding sites were located within the same TAD, as indicated by a higher contact frequency with the *SEMA7A* promoter. However, neither HSF1 binding site showed a local increase in contact frequency relative to the surrounding locus within the same TAD in either HS or NHS cells; nor did we observe any systematic difference in contact frequency with either HSF1 binding site when comparing NHS to HS.

Reasoning that we were likely underpowered to identify more subtle differences in contact frequencies at individual genes, we examined the entire set of HSF1-dependent HS up-regulated genes that have no evidence of HSF1 binding sites within 10 kb of their TSS (n = 49). We found no significant differences in contact frequency before and after HS (**Fig. 4C**). Examination of contact frequencies between HSF1-independent and down-regulated genes and the nearest active transcriptional regulatory element whose activity correlates with gene transcription based on PRO-seq, revealed no systematic differences in contact frequencies following HS at these gene classes (**Fig. 4C**). These results indicate that global changes in contact frequencies between enhancers and target genes remain largely unchanged following 30 min HS.

### Chromatin contacts established before HS accurately predict HSF1-dependent genes

A major unresolved problem in transcription regulation is identifying which enhancers regulate target genes. Having observed no substantial changes in contact frequency following HS that would allow us to connect target genes with HSF1 binding sites, we asked whether the chromatin contacts necessary to facilitate a robust HS response were established in the NHS condition. Consistent with this hypothesis, simply the distance to the nearest HSF1 binding site predicted genes that were dependent on HSF1 with reasonably high accuracy (**Fig. 5A, S4**). However, this criterion did not predict HSF1-dependent genes that were dependent on distal contacts, like *SEMA7A*.

**Figure 5:**
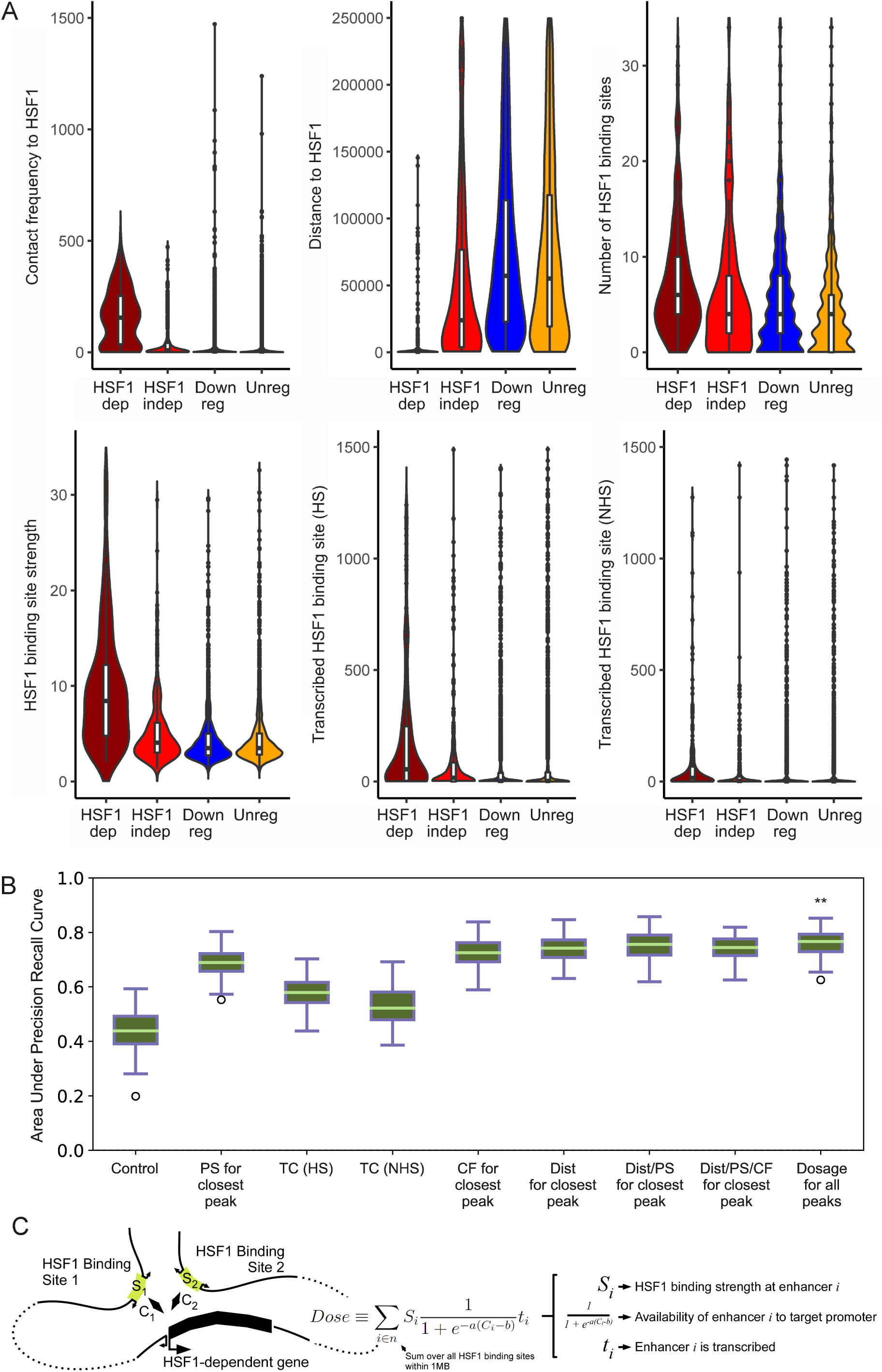
Prediction of HSF1-dependent up-regulated genes using NHS contact information. A) Comparison of the correlation between HSF1-dependent gene activation and various genomic data. Panels show: the NHS contact frequency between genes’ TSS and the closest HSF1 binding site (top left); the linear distance between genes’ TSS and the closest HSF1 binding site (top middle); the number of HSF1 binding sites within a 2 Mb window around each gene TSS (top right); the strength (fold enrichment of HSF1 signal) of the closest HSF1 binding site (bottom left), transcription count (PRO-seq reads) at the closest HSF1 binding site for HS data (bottom middle) and for NHS data (bottom right). B) Boxplots showing spread of area under the precision recall curve (auPRC) results for 1000 iterations of various classifiers attempting to distinguish HSF1-dependent genes from HSF1-independent genes. From left to right, **Control**: results obtained by randomly selecting the gene class; **PS for closest peak**: peak strength (fold enrichment of HSF1 binding signal) of the closest HSF1 binding site; **TC (HS)**: transcription count (PRO-seq reads) at closest HSF1 binding site for HS data; **TC (NHS)**: transcription count (PRO-seq reads) at closest HSF1 binding site for NHS data; **CF for closest peak**: contact frequency between genes’ TSS and the closest HSF1 binding site; **Dist for closest peak**: linear distance between genes’ TSS and the closest HSF1 binding site; **Dist/PS for closest peak**: linear distance / peak strength between genes’ TSS and the closest HSF1 binding site; **Dist/PS/CF for closest peak**: linear distance / peak strength / contact frequency between genes’ TSS and the closest HSF1 binding site; **Dosage for all peaks**: scaled contact frequency multiplied by peak strength for all HSF1 binding sites within a 2 Mb window around each gene TSS. C) Cartoon depicting components used to calculate HSF1 dosage.

Examination of the 49 HSF1-dependent genes that don’t have any detectable HSF1 binding within 10 kb of the TSS revealed that the majority of them still had HSF1 binding site(s) located within the same TAD (n= 35/ 49; 71%). However, we noticed no evidence of a focal increase in contact frequency between the nearest distal HSF1 binding site and the HSF1-dependent gene promoter in either heat condition, either at individual genes, or in aggregate. Increasing resolution to 3 kb by integrating published K562 Hi-C data likewise did not reveal any evidence for a focal loop connection between HSF1-dependent up-regulated promoters and the nearest HSF1 binding site (**Fig. S5**). Thus, despite evidence of a functional interaction between distal HSF1 binding sites and HSF1-dependent up-regulation of these genes, we observed no evidence of local increases in their contact frequencies.

We asked whether we could distinguish HSF1-dependent and -independent genes based on Hi-C contact frequencies, HSF1 binding location, and HSF1 binding strength. The number of HSF1 binding sites, the contact frequency between the HSF1 binding site and the promoter, HSF1 binding strength, and abundance of RNA polymerase at the HSF1 binding sites, were each correlated with whether genes were HSF1-dependent or not (**Fig. 5A**). Integrating these variables into a single classifier using gradient boosted trees distinguished HSF1-dependent genes from HSF1-independent up-regulated genes much more accurately than random guessing, as determined by the area under the precision recall curve (auPRC) on holdout sites not used during model training (**Fig. 5B and S6**). The best model used the distance between the promoter and the nearest HSF1 binding site, and the HSF1 binding strength, suggesting that simply distance and strength were enough to accurately classify most HSF1 dependent genes.

To develop a more biologically motivated classifier, we reasoned that HSF1 binding strength and the frequency of HSF1 binding site-promoter interactions were the two most important factors for a distal HSF1 binding site to regulate a target gene (22). We defined the “HSF1 dose” as the sum of all HSF1 binding sites within 1 Mb multiplied by their scaled contact frequency, and accounting for whether there is transcription at these HSF1 binding sites detectable by PRO-seq (23). We found that HSF1 dose predicted HSF1-dependent and -independent up-regulated genes slightly but significantly better than any other model (auPRC = 0.77; **Fig. 5B**). Notably, transcribed HSF1 binding sites had a larger effect on HSF1-dependent gene classification than non-transcribed enhancers, consistent with reports that many active enhancers are transcribed (24–26). Collectively, these results demonstrate that HSF1 dose (integrating HSF1 binding strength, transcription status, and contact frequency of nearby HSF1 binding sites) accurately predicted HSF1’s direct target genes.

### Pre-programmed chromatin architecture is conserved across metazoans

We asked whether chromatin architecture changes during heat stress in another metazoan organism. An earlier study with *Drosophila* Kc 167 cells has shown that TAD structures undergo reorganization upon heat shock with a general reduction in border strength (20). However, our results with heat shocked *Drosophila* S2 cells revealed no significant changes in TAD structure, insulation score or compartmentalization despite dramatic transcriptional activation in hundreds of genes (**Fig. 6A, B, and C**). Compartment calls and insulation scores in NHS and HS conditions were found to be highly similar, with both having a Pearson coefficient of 0.99 (**Fig. 6B and C**). The data recapitulated that observed in human K562 cells showing pre-established contacts between HSF dependent upregulated genes and their regulatory elements, in addition to showing a propensity towards a higher number of short range contacts when compared to the background contact frequency (**Fig. 6D**). We further analyzed the data published in NHS (27) and HS (20) conditions and found that technical variation between the NHS replicates could explain the some of the differences in chromatin conformation reported in (20) between NHS and HS cells (**Fig. S7**). Our analysis suggests that response to HS in the context of 3D genome organization is prewired across metazoans, and this could be necessary to provide the ‘power’ to rapidly drive the activation of genes having these preexisting connections.

**Figure 6:**
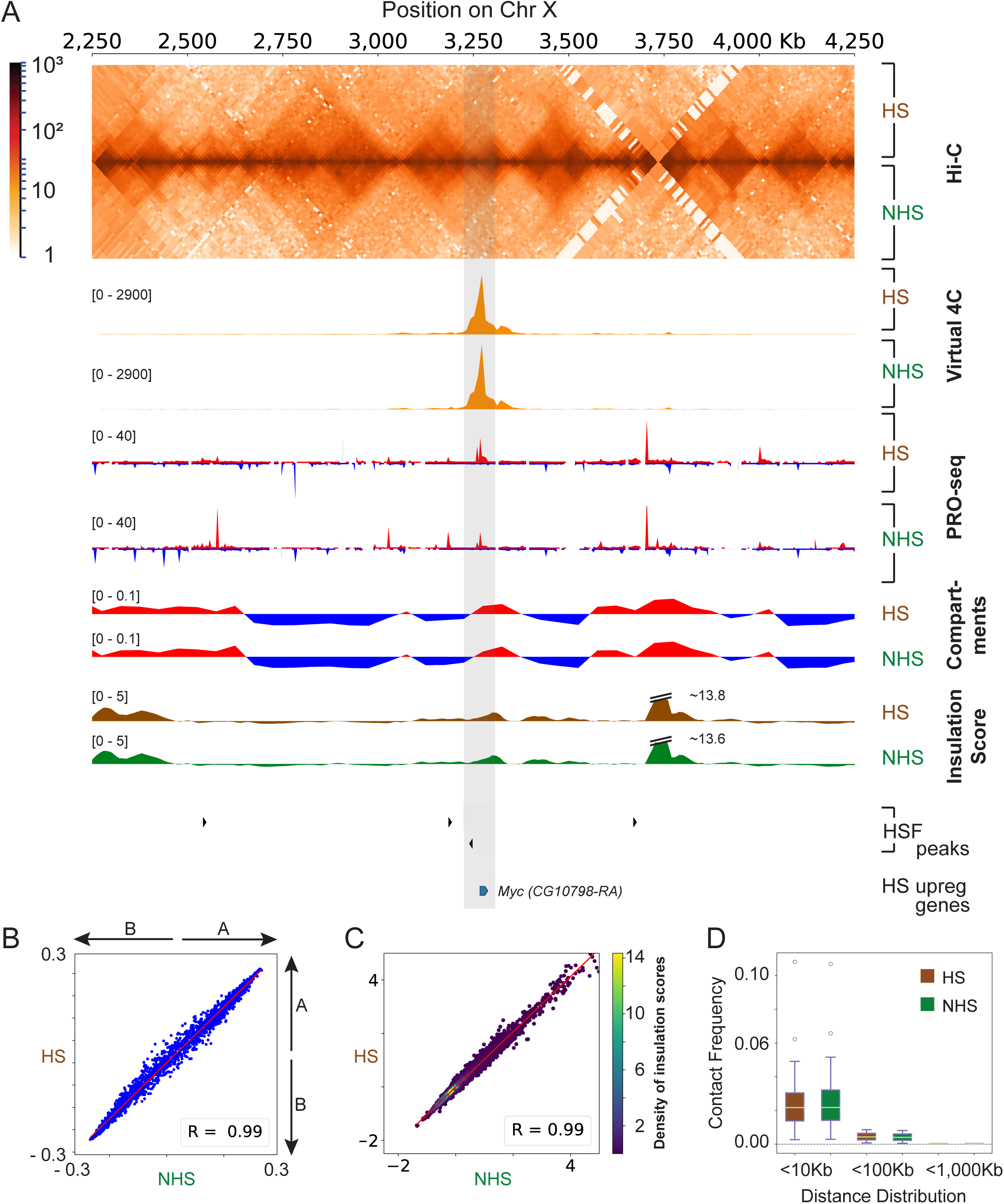
Chromatin conformations are highly similar under NHS and HS conditions in *Drosophila* S2 cells. A) Comparison of *in situ* Hi-C, Virtual 4C, PRO-seq, compartmentalization, and insulation scores for *Drosophila* S2 cells under NHS and HS conditions. B) Correlation between the strength of compartment calls before and after HS. Plots were drawn as in Fig. 2B. C) Correlation between insulation scores before and after HS. Plots were drawn as in Fig. 3B. D) Contact frequency vs distance distribution pairs for HS vs NHS datasets. Plots were drawn as in Fig. 4A.

## DISCUSSION

Chromosome conformation capture assays have provided powerful tools for interrogating chromatin contacts in specific cell types and conditions (28–32). Our understanding of the chromosome structure has been dramatically reshaped in the recent years by the identification of TADs, sub-TADs, loops, compartments and their roles in functional regulation of the genome in association with the architectural proteins (16, 18, 33-37). Generally, TADs are highly conserved across cell types, but they disappear along with compartments during mitosis (38). In differentiating cells, TADs are shown to be generally conserved, but disruption of their boundaries have been reported in cancers cells that could lead to oncogenesis (39). These studies indicate that in normal physiological circumstances cells do not undergo significant rearrangements in TAD structure. However, there is evidence that some loop changes and compartmental switching occur during cellular differentiation and senescence associated with changes in gene expression (19, 40, 41).

Responses towards different environmental signals vary depending on the nature of the signal, cell type and function. Transcriptional activation by heat shock is achieved by preconditioning the chromatin landscape of enhancers and promoters that allow establishment of promoter-proximal paused Pol II and the recruitment of critical transcription factors to release paused Pol II into productive elongation (8). In contrast, heat shock associated transcriptional repression of thousands of genes takes place by inhibiting paused Pol II release in mammals and reducing Pol II density along entire genes in flies (42). Our data suggests that heat shock does not alter TAD structures or weaken intra/inter-TAD boundaries in humans and flies. Additionally, we did not observe any significant switching or loss of compartmental strength following heat shock that is greater than that in our highly-correlated biological replicates **(Fig. S2).** Such an observation re-emphasizes that transcriptional response upon heat shock does not perturb global chromatin conformation, rather the changes in paused Pol II densities at TSSs or across gene bodies is achieved primarily by loss or recruitment of transcription factors and chromatin remodelers (15, 42).

Our findings contrast with the data and conclusions previously published for in *Drosophila* Kc167 cells (20). This study reported a reduction in TAD border strength and increase in inter-TAD interactions accompanied by redistribution of architectural proteins upon a 20 min HS. This led to an interesting and surprising model where disrupted TAD boundaries following a thermal stress allow Polycomb complex containing enhancer-promoter clusters that leads to gene repression. Such a dramatic reorganization model seems inconsistent with the evolutionarily conserved transcriptional down regulation seen of different ontological classes of genes across multiple species during HS (5, 6, 8). We did not analyze changes in Polycomb mediated long-range interactions upon thermal stress as it is beyond the scope of this study. However, if such interactions increase, they are not an effect of TAD reorganization as our data show no detectable changes in TAD structures upon HS.

Cellular state and physiology appear to be critical in determining the dynamics of enhancer-promoter or promoter-promoter interactions. Although it has been shown that regulatory contacts are newly formed or strengthened while cells are undergoing transcriptional changes (43–45), there is also evidence of pre-established enhancer-promoter interactions during stimuli activation, differentiation, development and stress (29, 43, 46, 47). This suggests that both dynamic and stable enhancer-promoter contacts could have contextual roles to regulate transcription of specific genes in a spatiotemporal manner. However, in the case of HS response we observed that contacts between HS regulated genes and the distal HSF1 bound regulatory element in the first 10 kb are preformed prior to HS. This result recapitulates the pre-established enhancer-promoter contacts as observed upon TNF-α stimulation of IMR90 cells (29) or during hypoxic stress in MCF-7 cells (47). Such evidence leads us to speculate that genomes have evolved to pre-wire not only the local chromatin architecture (8) but also the long-range regulatory interactions in 3D prior to stress so that the transcriptional response could be expedited. Cells are more frequently exposed to different kinds of stresses including thermal, osmotic, hypoxic etc. compared to other signaling cues for differentiation and development. It is possible that cells need an architectural platform to initiate immediate and rapid transcriptional response to survive thermal stress and a similar mechanism is likely to occur during other stresses as well. The similarity in the effects of heat shock on chromatin structure in humans and *Drosophila* suggests that stress response mechanism is evolutionarily conserved in terms of regulatory interactions.

We used HSF1 as a model system to explore how transcription factors regulate target genes at a distance. Using data from an HSF1 knockdown, we identified transcriptional changes that were dependent on HSF1, but without HSF1 binding nearby the TSS. We found that Hi-C data can help to distinguish genes where up-regulation following HS was dependent on HSF1 from those where it was not. The best model integrated the binding strength of HSF1 (as determined by ChIP-seq), the presence of eRNA transcription at an HSF1 binding site, and the Hi-C contact frequency. Our observations are consistent with the notion that Hi-C data is a surrogate for the degree to which a distal enhancer is in a position where it is able to regulate target promoters (although we note that simply the distance between the HSF1-dependent TSS and the nearest HSF1 binding site preformed nearly as well in our test). Furthermore, data from the NHS condition was just as informative as data from HS, and we observed no changes in contact frequency after HS. Taken together, our results suggest that chromatin contacts observed in Hi-C which are necessary for a robust HS are all in place prior to thermal stress.

Our study investigates an important connection of transcriptional regulation to chromatin interactions. In summary, we find the that chromatin interactions between regulatory elements and their target promoters appear to be prewired, and that the massive changes in transcription and chromatin following HS are not accompanied by significant changes in TADs, TAD boundaries, compartments, or looping interactions. Although we do not see a change in these chromatin conformations after a 30 min HS, when transcription has already changed dramatically at genes and enhancers, we cannot rule out that such conformational changes may occur at longer HS times. Given the current resolution of chromosome conformation capture techniques, we cannot comment on the changes in chromatin contacts at shorter distances that could have a role in the transcriptional response upon HS. Further analysis using new, higher-resolution chromatin conformation capture methodologies could allow delving more deeply into changes in chromatin structure and interactions triggered by the stress response.

## MATERIALS & METHODS

### Heat shock treatments & formaldehyde crosslinking

*Drosophila* S2 cells were grown in M3+BPYE medium with 10% FBS at 25**°**C. Cells were transferred to a shaking water bath maintained at room temperature for NHS or at 36.5**°**C for HS. Simultaneously, an equal volume of medium (without FBS), kept at room temperature or at 48**°**C was added into NHS or HS cells respectively. Cells were then incubated for 20 min. Cells were centrifuged at 500 x g at 4**°**C for 5 min. The medium was removed and cells were resuspended in 1x PBS. Crosslinking was done with the addition of formaldehyde to a final concentration of 1%, and incubated for 10 min at room temperature with occasional mixing. Glycine was added to a final concentration of 147 mM, and incubated at room temperature for 5 min with mixing. Cells were centrifuged and the supernatant was discarded. The cells were washed once with 1x PBS and the pellets were flash frozen in liquid nitrogen and stored at −80**°**C

Heat shock for human K562 chronic myelogenous leukemia cells were done according to the protocol described previously (8). The cells were grown in RPMI medium with 10% FBS at 37**°**C. Prior to treatment cells were concentrated in 10 ml medium. Cells were incubated at 37**°**C (NHS) or at 42**°**C (HS) water bath for 30 min. After treatment cells were centrifuged for 5 min at 4**°**C and the supernatant was discarded. Cells were resuspended in 1x PBS and crosslinked with formaldehyde to a final concentration of 1% for 10 min at room temperature with occasional mixing. Crosslinking was quenched by the addition of 200 mM glycine for 5 min at room temperature with mixing. Cells were centrifuged at 500 x g for 5 min at 4**°**C and the medium was discarded. The cells were washed once with 1x PBS and the pellets were flash frozen and stored at −80**°**C.

### In situ Hi-C

We used 58 million S2 or 5 million K562 cells for in situ Hi-C. The frozen crosslinked cell pellets were thawed on ice and resuspended in 250 μL ice-cold Hi-C lysis buffer (10mM Tris-HCl pH8.0, 10mM NaCl, 0.2% NP40) with Protease inhibitor cocktail (Thermo). Cells were incubated on ice for 30 min, centrifuged and washed once with 500 μL ice-cold Hi-C lysis buffer. The pellet was resuspended in 50 μL of 0.5% SDS and incubated at 62**°**C for 7 min followed by addition of 145 μL water and 25 μL of 10% Triton-X-100 and incubated at 37**°**C for 15 min. Finally, 25 μL of 10x NEBuffer2 and 5 μL (125 units) of MboI (NEB) was added to the mixture. The sample was digested at 37**°**C overnight with rotation. The sample was incubated at 62°C for 20 min to inactivate MboI and then cooled to room temperature. Biotin fill-in of digested ends was done by adding 1.5 μL each of 10 mM dCTP, dGTP, dTTP, 37.5 μL of 0.4 mM Biotin-14-dATP (Thermo) and 8 μL (40 units) of Klenow polymerase (NEB). The reaction was incubated at 37**°**C with rotation for 90 min with rotation. Ligation was performed upon addition of 100 μL 10% Triton-X-100, 120 μL of 10x T4 DNA ligase buffer (NEB), 12 μL of 10mg/ml BSA (NEB), 5 μL (2000 units) of T4 DNA ligase (NEB) and 663 μL of water. The ligation mixture was incubated at room temperature for 4 h with rotation. To reverse the crosslinks, 50 μL of 20 mg/ml Proteinase K (NEB) and 120 μL of 10% SDS were added to the sample followed by an incubation at 55**°**C for 30 min. To this mixture 130 μL of 5M NaCl was added and incubated at 68**°**C overnight. The reactions were cooled to room temperature and the sample was purified by 1.6x volume of 100% ethanol and 0.1x volume of 3M Sodium acetate, pH 5.2 followed by incubation at −80**°**C for 15 min. The sample was pelleted by spinning at 20,000 x g at 4**°**C and washed twice with 70% ethanol. The pellet was resuspended in 110 μL of 10 mM Tris-HCl, pH 8.0 and incubated at 37**°**C for 15 min to dissolve. The purified sample was sonicated using a Bioruptor Diagenode sonicator at low setting, with 30 second on and 90 second off for 20 min in an ice-cold water bath at 4**°**C. The sonicated sample was then treated with 2 μL RNase A/T1 cocktail (Thermo) for 30 min at 37**°**C. The DNA was cleaned up using Qiaquick PCR purification kit (Qiagen) and the eluate volume was brought up to 300 μL with 10mM Tris-HCl, pH 8.0. Biotin pull down was done with 150 μL of Dynabeads MyOne Streptavidin T1 (Thermo) that was washed with 400 μL of 1x Tween wash buffer (5 mM Tris-HCl, pH 7.5, 0.5 mM EDTA, 1M NaCl, 0.05% Tween-20). Washed beads were resuspended in 300 μL of 2x Binding and wash buffer (10 mM Tris-HCl, pH 7.5, 1 mM EDTA, 2 M NaCl) and added into 300 μL of DNA sample. Binding was done at room temperature for 15 min with rotation. Beads were washed twice with 1x Tween wash buffer, transferred into a new tube and heated at 55**°**C for 2 min with shaking in a thermomixer. Beads were then washed once with 100 μL of 1x NEB T4 DNA ligase buffer and transferred into a new tube. End repair was done by adding 88 μL of 1x NEB T4 DNA ligase buffer, 2 μL of 25 mM dNTP, 5 μL (50 units) of T4 PNK (NEB), 4 μL (12 units) of T4 DNA polymerase (NEB) and 1 μL (5 units) of Klenow polymerase (NEB). The mixture was incubated at room temperature for 30 min. Beads were washed with the 1x Tween wash buffer as described previously and once with 100 μL of 1x NEBuffer2, and transferred into a new tube. A-tailing was done by adding 90 μL of 1x NEBuffer2, 5 μL of 10 mM dATP and 5 μL (25 units) of Klenow (3-5’ exo-) polymerase (NEB). The reaction was incubated at 37**°**C for 30 min. Beads were washed with 1x Tween wash buffer just as before and once with 100 μL of 1x NEB T4 DNA ligase buffer, and transferred to a new tube. Adaptor ligation was done by adding 50 μL of 1.1x NEB T4 DNA ligase buffer, 2 μL of 3 μM Truseq/Universal indexed adaptor and 3 μL (1200 units) of T4 DNA ligase (NEB). Ligation was done at room temperature for 2 h. Beads were washed with 1x Tween wash buffer as described previously and test PCR was performed with P5 and P7 primers to determine the optimal cycle number for library amplification. Final PCR was done with 90% of the sample for 8-10 cycles. The final amplified library was purified with Qiaquick PCR purification kit (Qiagen) and sequenced using Illumina Nextseq 500.

### Hi-C data analysis

Files containing sequenced read pairs were processed using the Juicer pipeline as described (16). Reads for human K562 cells and *Drosophila* S2 cells were aligned to hg19 and dm3 respectively. We required that all alignments were high-quality by filtering for a MAPQ score greater than 30.

Reads for *Drosophila* Kc167 cells (Fig. S7) from (20) (available at Gene Expression Omnibus (GEO) database (http://www.ncbi.nlm.nih.gov/geo) under accession: GSE63518) were processed in a similar fashion (i.e. aligned to dm3 and processed using the Juicer pipeline).

Map resolution was calculated according to the definition proposed in ref (16), as the smallest bin size such that 80% of loci have at least 1,000 contacts. We used this definition to determine the finest scale at which one can reliably discern local features.

### Contact map similarity

Visualizations of the contact maps, PRO-seq data, and ChIP-seq data were produced using Juicebox (16), HiGlass (48), and pyGenomeTracks (21).

To measure contact map similarity beyond our initial visual comparison, we used HiCRep to obtain a stratum-adjusted correlation coefficient (SCC) (17). This was done for all chromosomes in human and *Drosophila*, in both cases using a bin size of 10Kb and maximum interaction distance set to 5Mb. For human, a smoothing parameter of 12 was used; for *Drosophila*, 8 was used, as recommended by the HiCRep software.

To determine compartment type (active or inactive) and compartmentalization strength, we computed the first principal component (PC) of the Pearson's correlation matrix of the observed contact map / expected contact map, across the entire genome. Pearson correlation matrices were computed using the Pearsons tool, and the PC computed using the eigenvector tool (16), both using a bin size of 50 Kb on KR normalized datasets. For *Drosophila* data, we used the “-p” option to the eigenvector tool to ignore sparsity when calculating the PC.

Typically, open / active compartment are defined as having positive PC1 values. However, we noted that the sign of the PC was reversed on some chromosomes. In order to confirm each compartment call, we used Pol II occupancy, as indicated by our PRO-seq data and the ChIP-seq signal for H3K36me3 (ENCODE) in the same region as markers for open chromatin to switch the PC sign when necessary. In all cases, we defined positive PC1 scores (the A compartment) as the compartment that correlated with active transcription and H3K36me3.

Insulation score was determined across the entire genome using the HiCExplorer hicFindTads tool (21). Default values were used for the ‘correctForMultipleTesting’ and ‘thresholdComparisons’ parameters (‘fdr’ and ‘0.01’ respectively), and ‘minDepth’, ‘maxDepth’, and ‘step’ were set to 30 Kb, 100 Kb, and 10Kb respectively.

We used gradient boosted trees implemented in scikit-learn’s ADABoostClassifier (49). Genes were randomly split into training and test datasets (80% of the data used for the training set and 20% held out for testing) and the classifier was trained on various combinations of features (or models) from this data (see below). All models were trained to discriminate between HSF1-dependent and HSF1-independent up-regulated genes on the basis of the nearest HSF1 binding site. A precision / recall curve was produced based on the correct and incorrect predictions made by the classifier and the area under the curve (AUC) calculated to assess of how well the classifier performed. This process of randomly splitting the data followed by training a classifier was performed 1000 times to get a spread of AUC values for each model. Values where plotted on a boxplot (Fig. 5B). A Wilcoxon rank-sum test was used to compare how each model performed relative to the others. Multiple models where tested in order to determine the predictive value of various features within our dataset. Results for the following models are shown in figure 5B:

Control: Bypasses the classifier predictions and represents the results obtained by random chance.

Dist / PS for closest peak: Uses the distance from the gene TSS to the closest peak and the strength of that peak.

Dist / PS / CF: Similar the previous model with the addition of the contact frequency to the closest peak.

We defined HSF1 dosage as:

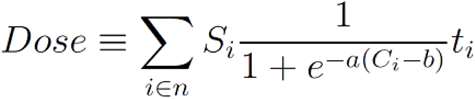

where *i* represents the HSF1 binding site from {1 .. n} that are within 1 megabase of each gene’s TSS; *Si* represents the fold enrichment of HSF1 signal in binding site *i* over input control called by MACS 1.4 (50); *ti* was set as a 1 if peak *i* intersected a dREG transcription initiation region (23) and otherwise was set to a value of *z* (see below). *C*_i_** was the contact frequency between HSF1 binding site *i* and the TSS; *a*, *b*, and *z* were free parameters of the model that were optimized to maximize auPRC on a training set of genes. In all cases, the training and test sets of genes were the same as used for the gradient boosted trees (described above). We noted that optimizing *a*, *b*, and *z* simultaneously using L-BFGS or conjugate gradients resulted in poor performance, in which the fitted values were nearly unchanged from the starting values. Therefore, we used Brent’s method (51) to identify the values of *a*, *b*, or *z* that maximized auPRC while holding the other values constant. We performed three rounds of optimization for each parameter. We bounded values of *a* = [0, 1], *b* = [0, 500], and *z* = [0,1].

### Additional datasets used in this study

K562 HSF1 ChIP-seq data - GSE43579 (7); K562 PRO-seq data - GSE89230 (8); K562 H4ac CHIP-seq data - GSE89382 (8); S2 HSF ChIP-seq data - GSE19025 (4); S2 PRO-seq data - GSE77607 (5).

### Code repository

All other code was custom written in Python 2.7. The significant parts of this code, and example data, are available on the Danko Lab‘s GitHub website (https://github.com/Danko-Lab/HS_transcription_regulation).

## ACKNOWLEDGEMENTS

We thank members of the Lis and Danko labs for thoughtful discussions about this work. CGD, AO, and JTL acknowledge support from the National Institutes of Health Common Fund 4D Nucleome Program (Grant U01HL129958). Additional support was provided by an NHGRI (National Human Genome Research Institute) R01-HG009309 and an NVIDIA GPU grant to CGD. The content is solely the responsibility of the authors and does not necessarily represent the official views of the US National Institutes of Health.

---

**Figure S1:**
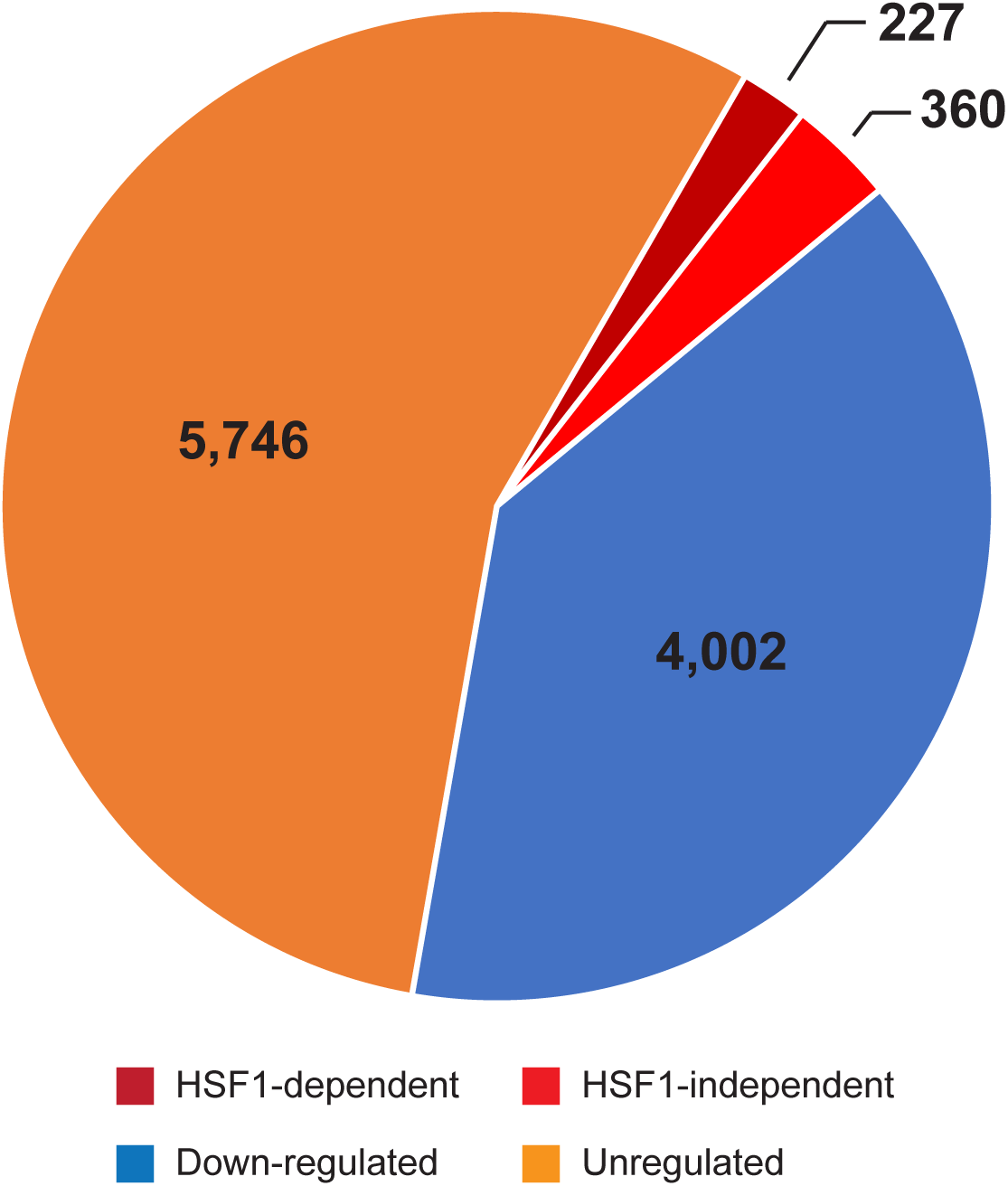
Distribution of genes between various classes (HSF1-dependent and HSF1-independent up-regulated, HS down-regulated, and HS unregulated).

**Figure S2:**
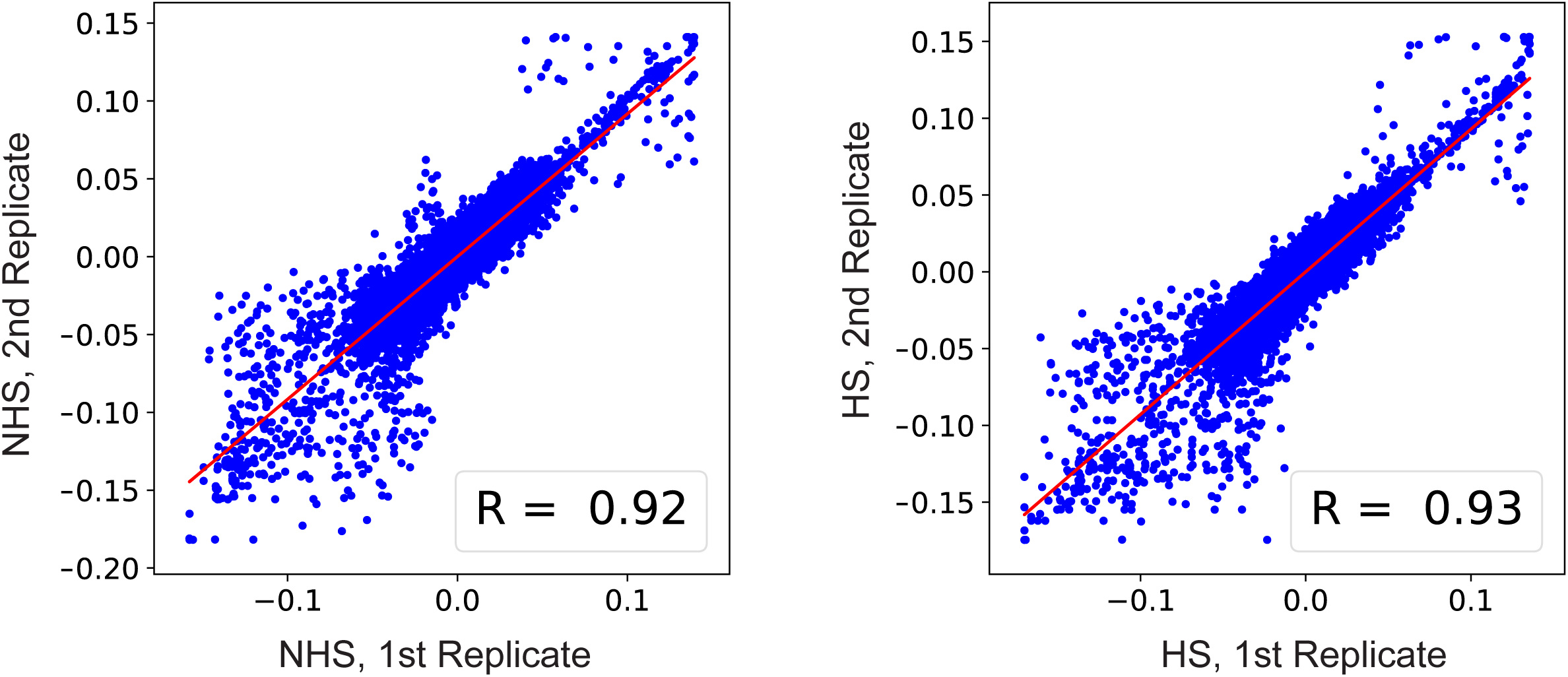
Correlation between the strength of compartment calls for replicates from the same condition, genome-wide in K562.

**Figure S3:**
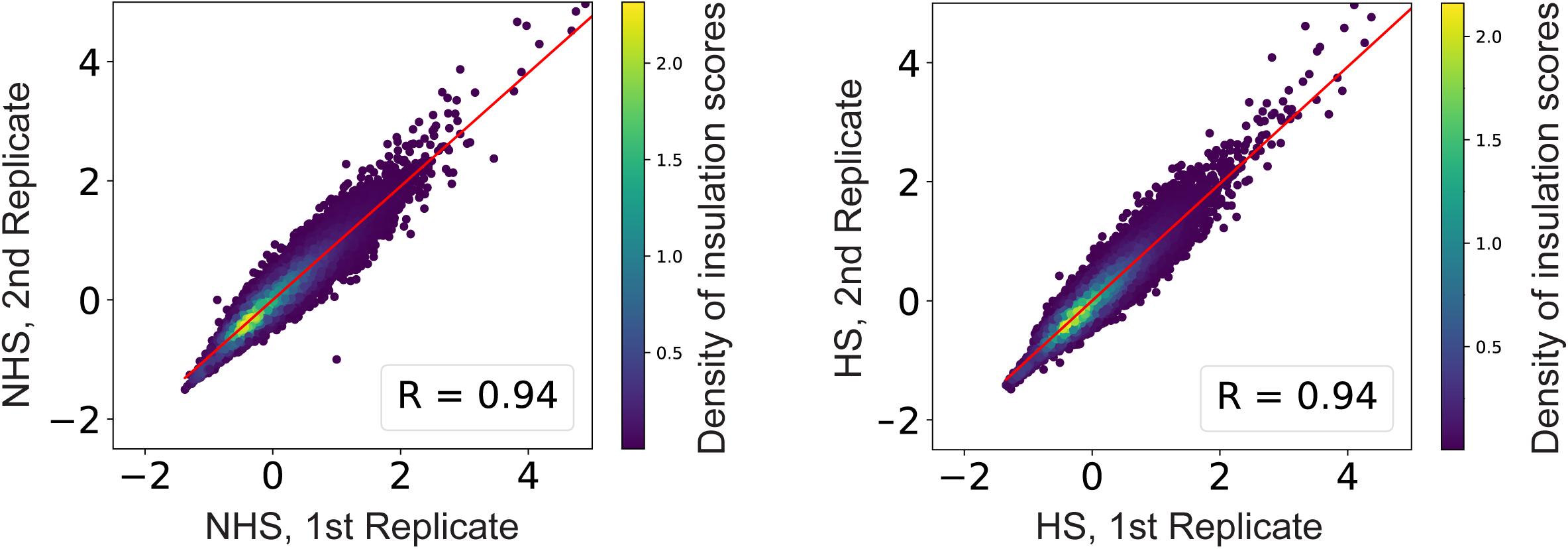
Correlation of insulation scores for replicates from the same condition, genome-wide in K562.

**Figure S4:**
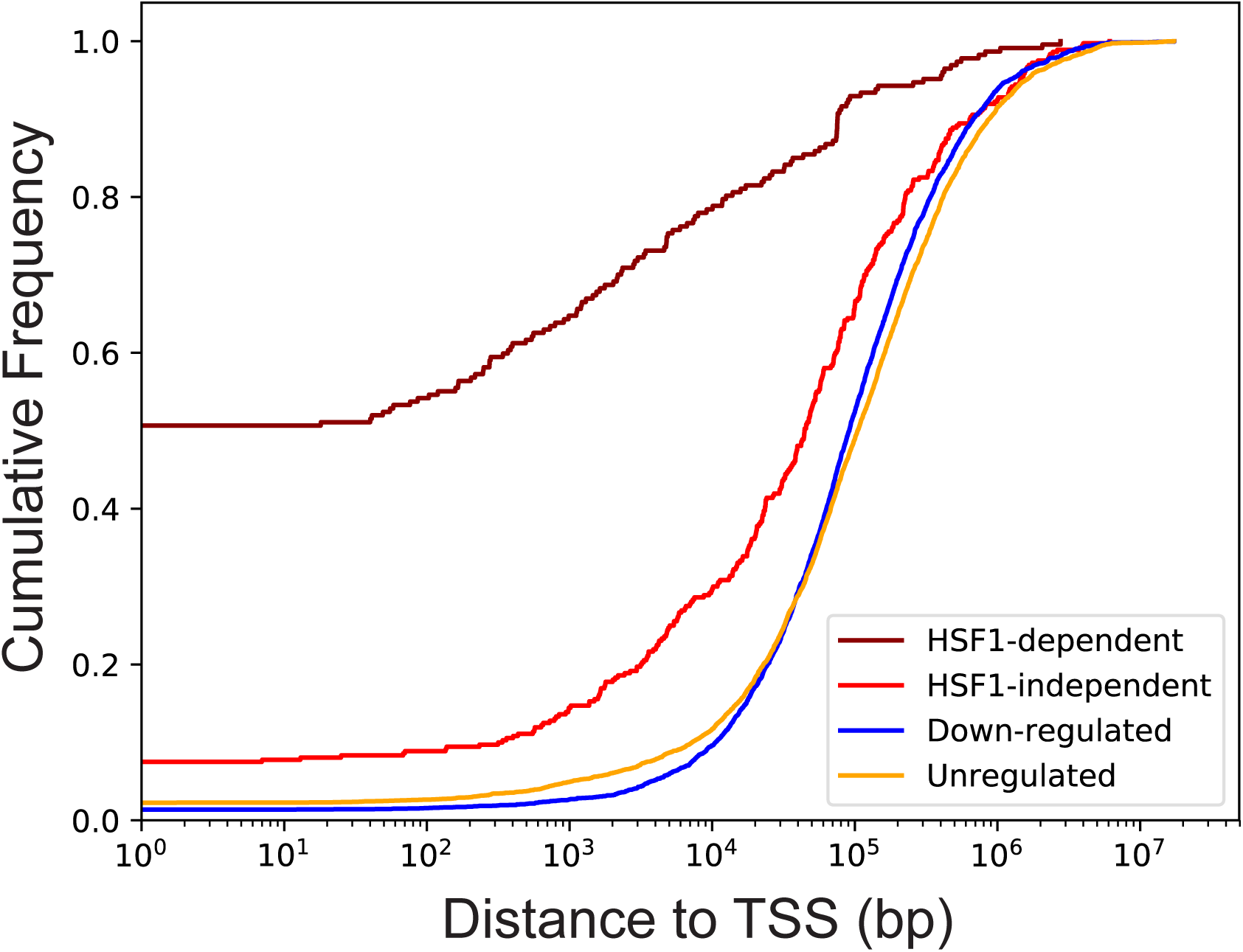
Cumulative frequency of the distances in base pairs between the peaks of HSF1 binding sites and TSS of HSF1-dependent, HSF1-independent, down-regulated and unchanged genes.

**Figure S5:**
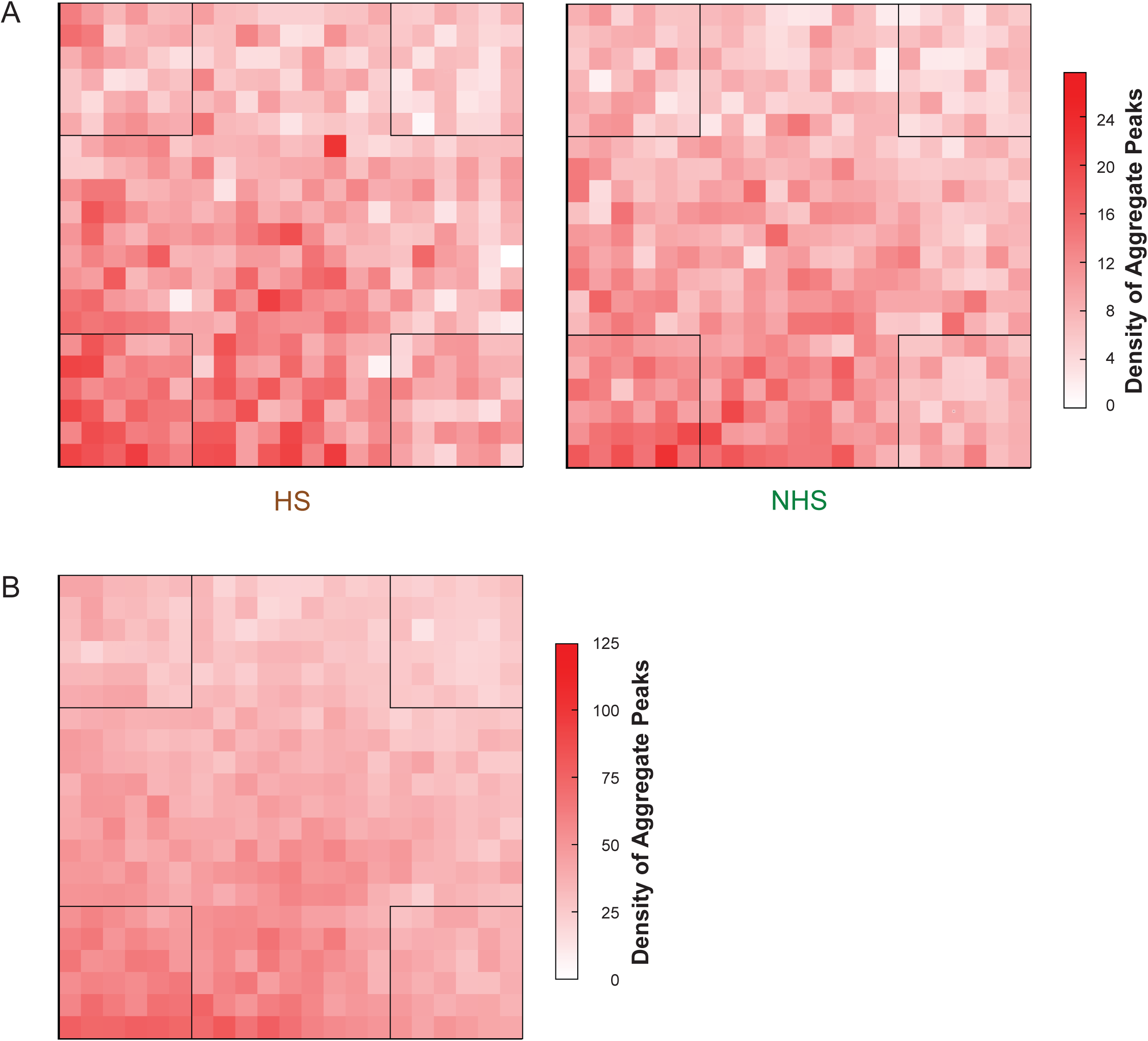
Aggregate peak analysis for contact frequency between the nearest distal HSF1 binding site and the HSF1-dependent gene TSS A) No evidence of a focal increase in contact frequency between the nearest distal HSF1 binding site and the HSF1 dependent gene was observed in either heat condition. Aggregate peak analysis was done as described previously (16). B) K562 Hi-C data from (16) integrated with data from this study, increasing resolution to 3 kb.

**Figure S6:**
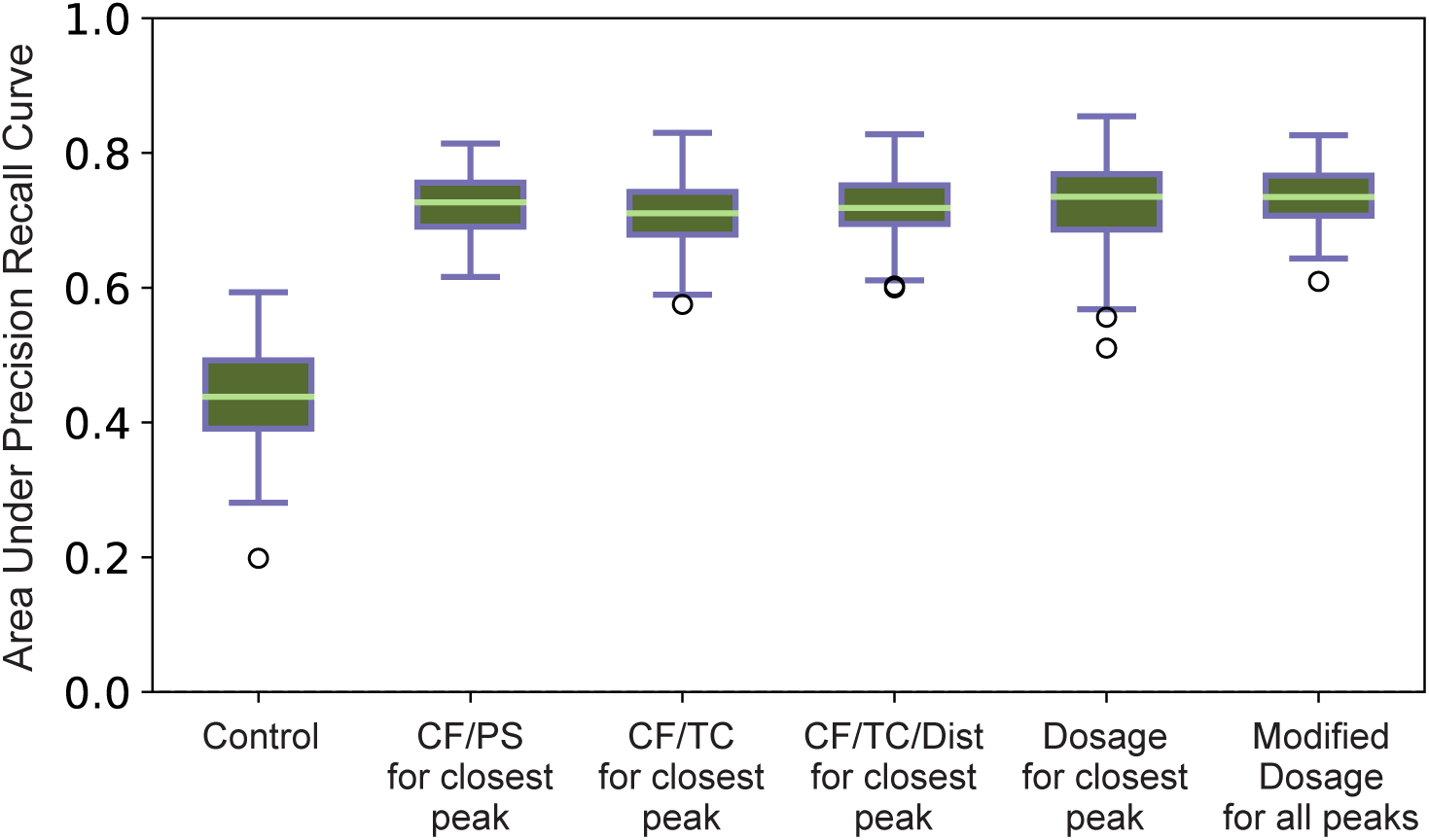
Boxplots showing spread of area under the precision recall curve (auPRC) results for 1000 iterations of classifiers not included in Fig. 5B. From left to right, **Control**: results obtained by randomly selecting the gene class; **CF/PS for closest peak**: contact frequency and peak strength (fold enrichment of HSF1 binding signal) of the closest HSF1 binding site; **CF/TC for closest peak**: contact frequency and transcribed count (PRO-seq reads) at the closest HSF1 binding site for NHS data; **CF/TC/Dist for closest peak**: contact frequency, linear distance, and transcribed count (PRO-seq reads) at the closest HSF1 binding site for NHS data; **Dosage for closest peak**: scaled contact frequency multiplied by peak strength for closest HSF1 binding site; **Modified dosage for all peaks**: scaled contact frequency multiplied by peak strength for all transcribed HSF1 binding sites within a 2 Mb window around each gene TSS.

**Figure S7:**
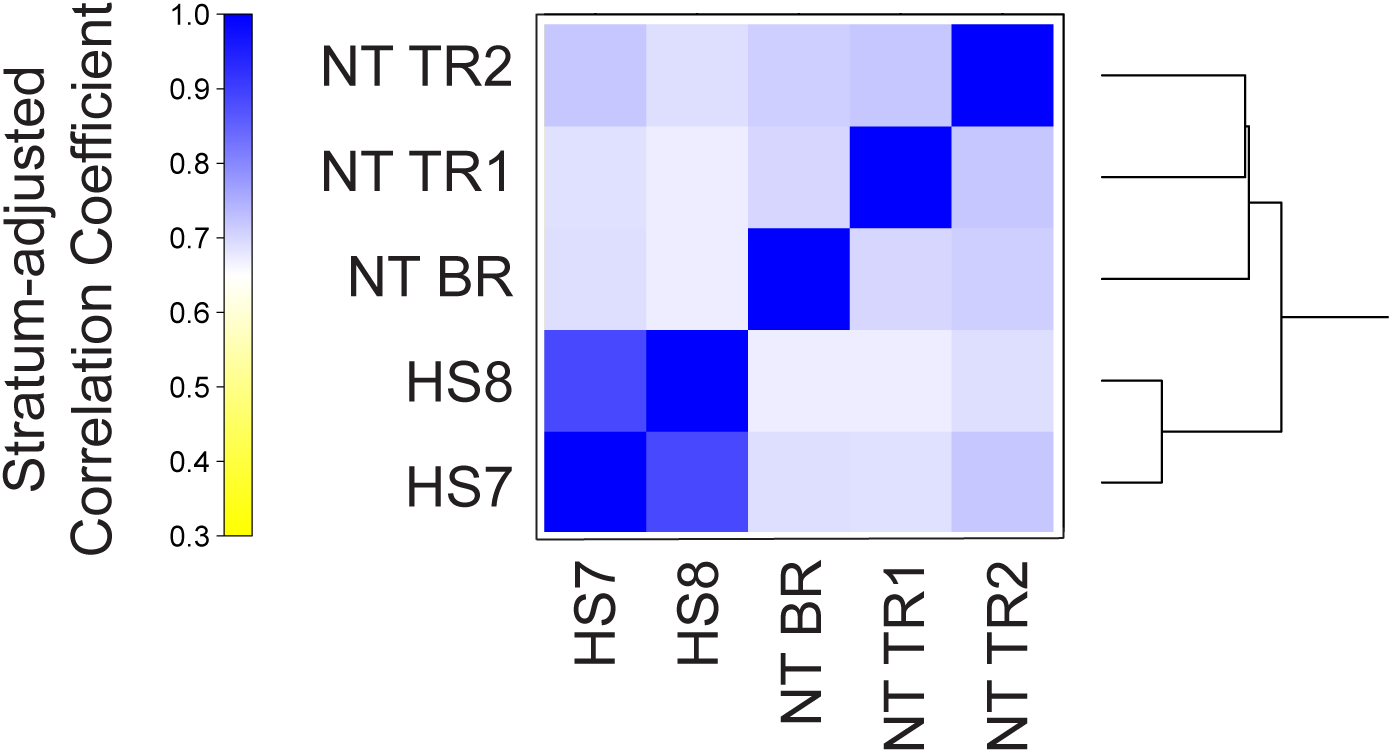
Stratum-adjusted correlation coefficients for heat shocked and non-treated (NT/NHS) replicates. The highest-level separation is between the HS (20) and non-treated conditions (27), although this could be a technical variation between datasets. The next highest level of separation is between each of the non-treated replicates.

**Table S1.**
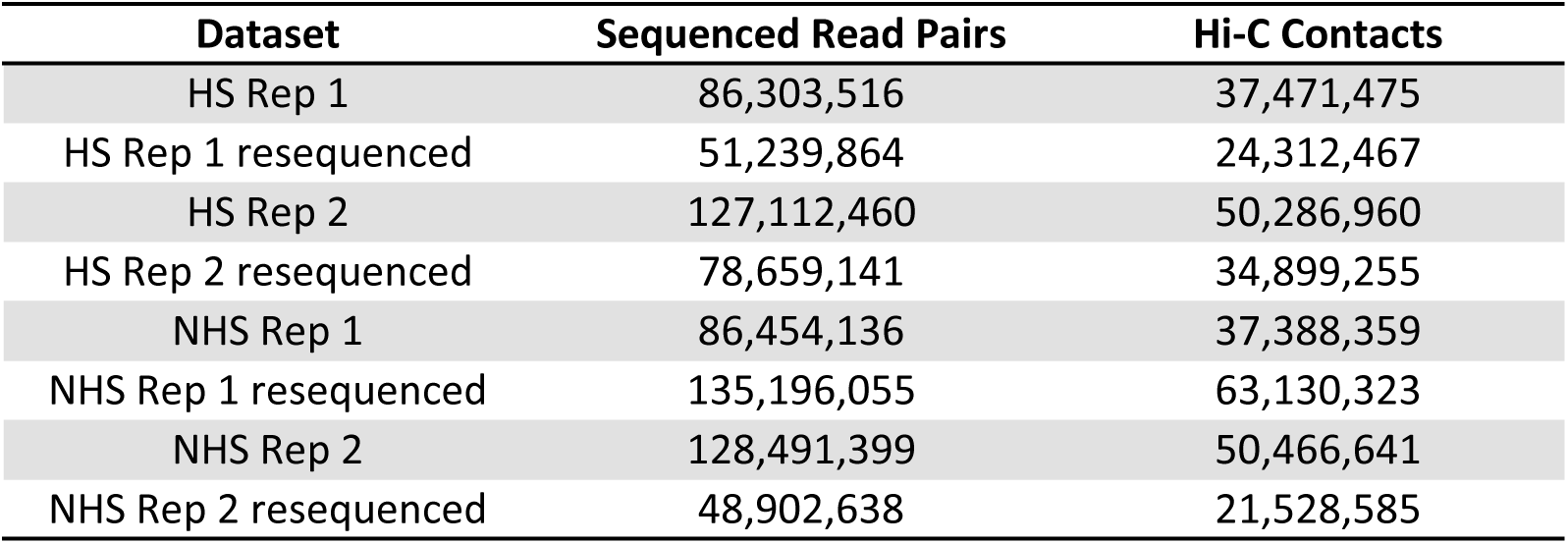

| Dataset | Sequenced Read Pairs | Hi-C Contacts |
| --- | --- | --- |
| HS Rep 1 | 86,303,516 | 37,471,475 |
| HS Rep 1 resequenced | 51,239,864 | 24,312,467 |
| HS Rep 2 | 127,112,460 | 50,286,960 |
| HS Rep 2 resequenced | 78,659,141 | 34,899,255 |
| NHS Rep 1 | 86,454,136 | 37,388,359 |
| NHS Rep 1 resequenced | 135,196,055 | 63,130,323 |
| NHS Rep 2 | 128,491,399 | 50,466,641 |
| NHS Rep 2 resequenced | 48,902,638 | 21,528,585 |

**Table S2.**

| K562 Datasets | Stratum-adjusted Correlation Coefficient |
| --- | --- |
| HS 1st rep vs. HS 2nd rep | 0.905 |
| HS 1st rep vs. NHS 1st rep | 0.905 |
| HS 1st rep vs. NHS 2nd rep | 0.896 |
| HS 2nd rep vs. NHS 1st rep | 0.899 |
| HS 2nd rep vs. NHS 2nd rep | 0.915 |
| NHS 1st rep vs. NHS 2nd rep | 0.898 |

| S2 Datasets | Stratum-adjusted Correlation Coefficient |
| --- | --- |
| HS 1st rep vs. HS 2nd rep | 0.943 |
| HS 1st rep vs. NHS 1st rep | 0.955 |
| HS 1st rep vs. NHS 2nd rep | 0.928 |
| HS 2nd rep vs. NHS 1st rep | 0.971 |
| HS 2nd rep vs. NHS 2nd rep | 0.979 |
| NHS 1st rep vs. NHS 2nd rep | 0.972 |

